# Comprehensive Molecular Epidemiology of Influenza Viruses in Brazil: Insights from a Nationwide Analysis

**DOI:** 10.1101/2024.07.04.602044

**Authors:** Isabela Carvalho Brcko, Vinicius Carius de Souza, Gabriela Ribeiro, Alex Ranieri Jeronimo Lima, Antonio Jorge Martins, Claudia Renata dos Santos Barros, Eneas de Carvalho, James Siqueira Pereira, Loyze Paola Oliveira de Lima, Vincent Louis Viala, Simone Kashima, Debora Glenda Lima de La-Roque, Elaine Vieira Santos, Evandra Strazza Rodrigues, Juliana Almeida Nunes, Leandro Spalato Torres, Luiz Artur Vieira Caldeira, Melissa Palmieri, Caio Genovez Medina, Raphael Augusto de Arruda, Renata Beividas Lopes, Geraldo Reple Sobrinho, Daniel Macedo de Melo Jorge, Eurico Arruda, Eladja Christina Bezerra da Silva Mendes, Hazerral de Oliveira Santos, Arabela Leal e Silva de Mello, Felicidade Mota Pereira, Marcela Kelly Astete Gómez, Vanessa Brandão Nardy, Brenno Vinicius Martins Henrique, Lucas Luiz Vieira, Mariana Matos Roll, Elaine Cristina de Oliveira, Júlia Deffune Profeta Cidin Almeida, Stephanni Figueiredo da Silva, Gleissy Adriane Lima Borges, Katia Cristina de Lima Furtado, Patricia Miriam Sayuri Sato Barros da Costa, Shirley Moreira da Silva Chagas, Esper G. Kallás, Daniel Larh, Marta Giovanetti, Svetoslav Nanev Slavov, Sandra Coccuzzo Sampaio, Maria Carolina Elias Sabbaga

## Abstract

Influenza A and B viruses pose significant global health threats, with substantial impacts on morbidity and mortality. Understanding their molecular epidemiology in Brazil, a key hub for the circulation and dissemination of these viruses in South America, remains limited. This study, part of the Center for Viral Surveillance and Serological Assessment (CeVIVAS) project, addresses this by analyzing data and samples from all Brazilian macroregions, along with publicly available sequences from 2021-2023.

Phylogenetic analysis of the Hemagglutinin (HA) segment of Influenza A/H1N1pdm09, A/H3N2, and Influenza B/Victoria-lineage revealed the predominance of A/H3N2 2a.3 strain in 2021 and early 2022. This was succeeded by A/H3N2 2b until October 2022, after which A/H1N1pdm09 5a.2a and 5a.2a.1 lineages became prevalent, maintaining this status throughout 2023. B/Victoria circulated at low levels between December 2021 and September 2022, becoming co-prevalent with A/H1N1pdm09 5a.2a and 5a.2a.1 lineages.

Comparing the vaccine strain A/Darwin/9/2021 with circulating A/H3N2 viruses from 2021-2023 revealed shared mutations to aspartic acid at residues 186 and 225, altering the RBD domain’s charge. For A/H1N1pdm09, the 2022 consensus of 5a.2a.1 and the vaccine strain A/Victoria/2570/2019 had 14 amino acid substitutions. Key residues such as H180, D187, K219, R223, E224, and T133 are involved in hydrogen interactions with sialic acids, while N130, K142, and D222 may influence distance interactions based on docking analyses.

Distinct Influenza A lineage frequency patterns across Brazil’s macroregions underscore regional variations in virus circulation. This study characterizes the dynamics of Influenza A and B viruses in Brazil, offering valuable insights into their circulation patterns. These findings have significant public health implications, informing strategies to mitigate transmission risks, optimize vaccination efforts, and enhance outbreak control measures.

**Author summary:** This study investigates the molecular epidemiology of Influenza A and B viruses in Brazil from 2021 to 2023. Utilizing data from the Center for Viral Surveillance and Serological Assessment (CeVIVAS) and public databases, we performed a comprehensive phylogenetic analysis of the Hemagglutinin segments of Influenza A/H1N1pdm09, A/H3N2, and B/Victoria-lineage viruses across all Brazilian macroregions. Key findings reveal that the A/H3N2 2a.3 strain was predominant in 2021 and early 2022, followed by A/H3N2 2b, and later by A/H1N1pdm09 5a.2a and 5a.2a.1 lineages in late 2022 and throughout 2023. The B/Victoria strain circulated at low levels initially and later co-prevailed with A/H1N1pdm09 lineages. Comparing the vaccine strain A/Darwin/9/2021 with circulating A/H3N2 viruses from 2021-2023 and A/Victoria/2570/2019 with 5a.2a.1 of A/H1N1pdm09 circulating in 2022 revealed significant mutations which could affect the interaction of the viruses with sialic acids and potentially impact vaccine efficacy. Notably, we identified a substitution pattern among the predominant Influenza subtypes and observed distinct regional variations in Influenza A lineage frequencies across Brazil. These findings are critical for optimizing vaccination strategies and provide valuable data to inform public health policy and improve health outcomes.

## Introduction

Influenza A and B viruses cause annual worldwide epidemics, significantly impacting public health due to their high morbidity and mortality rates [1, 2]. These infections predominantly manifest as respiratory illnesses, exhibiting varying severity and serious complications, particularly among vulnerable demographics such as children and the elderly [3–5]. Annually, Influenza is responsible for over 200,000 deaths globally [6–8].

In Brazil, as well as globally, the primary causative agents of seasonal Influenza are Influenza A subtypes, particularly H1N1 and H3N2 [9, 10]. However, since 2001, there has been a co-circulation of Influenza B lineages, Victoria and Yamagata, with the Victoria lineage emerging as the predominant strain in recent seasons [11, 12].

Previous studies have indicated significant variation in the evolutionary patterns of Influenza A and B viruses based on viral strain [13, 14]. While Influenza A/H3N2 subtypes exhibit relatively rapid evolution with frequent replacements occurring every 2 to 5 years, A/H1N1pdm09 and Influenza B viruses evolve more slowly [15]. Despite this slower evolution, the co-circulation of multiple lineages allows for the emergence of new antigenic variants approximately every 3–8 years [16, 17].

Furthermore, these studies underscore the critical role of genomic surveillance of Influenza for understanding its epidemiology, incidence, and phylogenetic relationships, which may impact disease severity [18, 19]. Hemagglutinin, the primary antigenic target of Influenza vaccines, plays a crucial role in the pathogenesis of the infection and is subject to antigenic changes that can affect vaccine efficacy [10, 20, 21].

Despite Brazil’s established protocol for Influenza sample collection and vaccine strain selection [9], there remain gaps in the comprehensive genetic and phylogenetic characterization of Influenza A and B viruses in the country. This study aims to address these gaps by assessing and characterizing Influenza A and B viruses collected across the five Brazilian macroregions between 2021 and 2023 through phylogenetic analysis of the HA gene.

## Materials and methods

### CeVIVAS Dataset

#### Clinical Sample Collection

The CeVIVAS project, a collaborative initiative involving the Instituto Butantan, Hemocentro Ribeirão Preto, several Central Public Health Laboratories (LACEN) across Brazil, and the municipalities of São Paulo and São Bernardo do Campo, aims to enhance viral surveillance and serological assessment. To ensure representative sampling of Influenza, CeVIVAS included LACENs from all five Brazilian macroregions: Paŕa (North), Alagoas and Bahia (Northeast), Federal District (Braśılia), Mato Grosso (Midwest), Ribeirão Preto, São Paulo, and São Bernardo do Campo cities (Southeast), and Parańa (South).

Only samples that tested positive for Influenza viruses with Ct values ¡ 30 accompanied by available epidemiological metadata, such as the date and sample collection location, were selected and sequenced. This stringent criterion ensures the inclusion of informative samples for a more comprehensive assessment of the prevalence and genetic diversity of circulating Influenza A and B viruses in Brazil.

This comprehensive collection effort yielded 1,277 newly generated Brazilian HA sequences, distributed as follows: 313 samples of Influenza A/H1N1pdm09: 56 from Alagoas, 4 from Bahia, 50 from the Federal District, 23 from Mato Grosso, and 180 from São Paulo; 713 samples of Influenza A/H3N2, with contributions from Alagoas (126), Bahia (235), Paŕa (22), the Federal District (86), Mato Grosso (69), and São Paulo (175), and 252 samples of Influenza B/Victoria lineage, 130 from Alagoas and 122 from São Paulo. It is noteworthy that all sequences, but one, exhibit a minimal 83% coverage and complete HA CDS gene. For further details, refer to S5 Fig and S8 Table. All sequences generated in this study were deposited in the GISAID database.

The sampling plan was meticulously designed to ensure precision and representativeness of the collected data. We utilized a margin of error (alpha error) of 5% and a confidence interval (CI) of 95%, allowing for precise estimation of population parameters. Additionally, we established a sampling power of at least 80%, ensuring the study’s ability to detect true differences or effects if present in the population. The final calculated sample size was increased by 20% to account for possible losses.

Furthermore, acknowledging the influence of seasonality on influenza prevalence, we adjusted the sample to seasonal variation. During high prevalence months (October to May), we reduced the sample by 0.25, and in the remaining months, we increased it by 0.5 [22]. This approach ensured adequate sample size in each phase of the study, maintaining the representativeness of the collected data. The sample was calculated at the state level to guarantee precision and representativeness in each macroregion. Furthermore, the sample selection involved municipal diversity within each state.

#### Influenza A and B Whole Genome Sequencing

The positive clinical samples for Influenza A and B were extracted following the methodology described in Supporting information. Influenza A genomic sequences were obtained using universal primers for Influenza A (Opti1-F1, Opti1-F2, and Opti1-R1) as previously described by Mena et al. (2016), with minor modifications, for specific details on the reaction conditions refer to Supporting information. Influenza B genomes were obtained using the same one-step amplification strategy however utilizing a set of Influenza B universal primers described by [23]. Pooled libraries were sequenced using 2x150 bp pair-end flow cell kits for NextSeq2000 or 2x150 bp pair-end flow cell kits for MiSeq (Illumina). For more information about genomic library preparation and Influenza next-generation sequencing see Supporting information.

#### Genome Assembly Pipeline

The raw reads were submitted to an assembly pipeline consisting of three main steps, as depicted in S6 Fig. Initially, Trimmomatic was employed to exclude low-quality reads, adapters, and primer sequences [24].

Subsequently, the filtered reads underwent the first main step of the pipeline, where VAPOR [25] and an Influenza genome database were utilized. The Influenza genome database comprises sequences sourced from GISAID and GenBank, provided by INSaFLU [26]. The objective was to identify the best reference sequence for each segment of the Influenza virus. To achieve this, the segment sequence with the highest score was extracted from the database using seqtk [27] (available at: https://github.com/lh3/seqtk). This selected reference segment was then employed in the second main step, which involved individual assembly of each genomic viral segment.

The filtered reads were mapped to the selected reference segment using Bowtie2, with the –very-sensitive parameter for segment-specific reads selection [28]. Properly paired reads were extracted using SAMtools and BEDtools [29, 30]. The segment-specific reads were subsequently submitted to assembly using SPAdes [31]. The refinement process entailed mapping the scaffolds to the selected segment reference using Minimap2 with default parameters [32]. A pre-consensus was then generated using SAM tools and iVar [33]. An in-house Python script was applied to substitute degenerated bases inserted by iVar for Ns, as the assembled scaffolds may contain potential SNPs.

To obtain the final polished consensus segment, the filtered reads were mapped against the pre-consensus using Bowtie2. The iVar tool was employed with the parameters -m 10 and -q 20, requiring a minimum depth of 10 reads and a frequency of 25%. All the assembled segments from the sample were combined to generate a final genome fasta file. In cases where segment 4, which codes for the HA gene, was successfully assembled, it was subjected to clade attribution using Nextclade [34]. The clade attribution was performed by comparing the assembled segment against the following references: H1- A/Wisconsin/588/2019 (MW626062), H3- A/Darwin/6/2021 (EPI1857216), Victoria- B/Brisbane/60/2008 (KX058884), Yamagata - B/Wisconsin/01/2010 (JN993010).

### GISAID Dataset

To expand our dataset, we obtained additional HA gene sequences from the Global Initiative on Sharing All Influenza (GISAID) EpiFlu Database [35] (available at: https://www.gisaid.org/, accessed September 2023). This database includes sequences from Brazil and worldwide, from 2021 to 2023. We concentrated our analysis on the Influenza B/Victoria lineage and the Influenza A virus subtypes A/H3N2 and A/H1N1pdm09, given the absence of the Influenza B/Yamagata lineage detections since 2019 [11]. The dataset from Brazil, encompassing all macroregions, included 604 HA sequences for Influenza A/H1N1pdm09, 1,637 for A/H3N2, and 527 for B/Victoria. To perform a comprehensive phylogenetic analysis and minimize bias we sample non-Brazilian sequences across regions, including Argentina, Australia, China, South Africa, the United States, and the United Kingdom (see S10 Table). These locations were carefully chosen to encompass the world’s major geographic subdivisions and to ensure the availability of deposited sequences. Argentina, Australia, and South Africa were selected to represent countries in the Southern Hemisphere, spanning the continents of South America, Oceania, and Africa, respectively. Conversely, China, the United States, and the United Kingdom were chosen to represent countries in the Northern Hemisphere covering Asia, North America, and Europe, respectively [36]. We selected only high-quality sequences, with at least 85% coverage and complete HA CDS gene. For detailed information, see S8 Table.

Also, we selected hemagglutinin (HA) sequences from WHO-recommended trivalent egg-based vaccine strains for the 2021 to 2024 epidemic seasons in the Southern Hemisphere (SH) [37–39]. Specifically, for the Influenza A/H1N1pdm09 dataset, the strains A/Victoria/2570/2019, A/Sydney/5/2021, and A/Victoria/4897/2022 were included. For the Influenza A/H3N2 dataset, we added A/Hong Kong/2671/2019, A/Darwin/9/2021, and A/Thailand/8/2022. And in the dataset for the Influenza B/Victoria lineage, we included B/Washington/02/2019 and B/Austria/1359417/2021. Detailed information about these vaccine strains can be found in S11 Table.

For our final dataset, we merged CeVIVAS, GISAID, and Vaccine datasets, resulting in 3,966, 8,609, and 3,996 sequences from A/H1N1pdm09, H3N2, and B/Victoria-lineage, respectively. For more information see S8 Table.

### Phylogenetic Analysis

The phylogenetic trees of the A/H1N1pdm09, A/H3N2, and Influenza B/Victoria were inferred using the Nextstrain pipeline [34] (build available in https://github.com/nextstrain/seasonal-flu). Clades and subclades are assigned using the collection of signature mutations provided by Nextstrain for each lineage of A/H1N1pdm09, A/H3N2, and B/Victoria. The maximum-likelihood method using IQTree [40], was applied under the general time-reversible (GTR) model. To statistically support the phylogenetic trees, we applied ultrafast Bootstrap (UFBoot) using 1000 replicates [40]. UFBoot confidence values *>*70% were considered as the cut-off for clustering. In the Nextstrain pipeline, during phylogenetic analysis and refining steps, samples that significantly deviate from the molecular clock model are automatically removed, resulting in phylogenetic trees comprising 3,517, 8,420, and 3,194 sequences from A/H1N1pdm09, A/H3N2, and B/Victoria-lineage, respectively. All trees were edited in R using the ggtree package [41].

### Genetic Analysis of Brazilian Influenza Lineages

#### Comparison of Vaccine Strains and Brazilian Consensus

To identify the presence of amino acid substitution in the HA segment in Brazilian sequences, we first categorized them based on their subclade/lineage and year of circulation. Next, only complete sequences with 100% coverage were selected, and a consensus sequence was generated by selecting the bases that appeared in at least 50% of the circulating sequences for each subclade/year group. We then compared the consensus sequences obtained from the Brazilian sequences with the vaccine strains recommended by the WHO for the Southern Hemisphere in the seasons spanning from 2021 to 2024. These comparisons were performed using amino acid sequences. The HA epitope regions were defined according to the Nextstrain pipeline, which is a widely used tool for the analysis of viral genetic sequences [42]. This approach aimed to identify any potential amino acid substitutions in the HA segment of Brazilian sequences compared to the vaccine strains, providing insights into the antigenic variability and potential implications for vaccine efficacy.

#### 0.0.1 Protein Structural Prediction Analysis

The protein structure prediction was performed using Modeller software version 10.4 [43]. To predict the structure of A/H1N1pmd09 hemagglutinin, we used the PDB6UYN, PDB6HJQ, and PDB5C0S structures as templates. For A/H3N2 hemagglutinin, we used the PDB4O58 structure as a template. All templates were selected prioritizing identity (*>*80%), coverage (*>*80%), resolution (*<*3.0Å), and their interaction with antibodies. After predicting the structures, we performed protonation using PDB2PQR v3.0 [44]. This step adjusts the ionization states of amino acid residues based on the input data, such as the pH value. We used the Propka method to predict the titration state of amino acid residues at pH 7.4 [45, 46]. To perform all electrostatic analyses, we used the APBS v5.0 software (Jurrus et al., 2018). For molecular docking assays, we used the Autodock Vina program [47]. We prepared the vaccine strain egg-propagated A/Victoria/2570/2019 and the consensus of subclade 5a.2a.1 from 2022, as well as the sialic acid (alpha-2,3, and alpha-2,6) at pH 7.4. The Gasteiger partial charge assignment was performed using the MGLTools 1.5.6 program (available at: https://ccsb.scripps.edu/mgltools). The grid center for molecular docking was set at the RDB domain, with the coordinates x=-17.345, y=-75.700, and z=48.168. The dimensions of the grid were set as X=18Å, Y=22Å, Z=18Å. This grid represents the region where the docking simulations will be performed to predict the binding affinity between the HA protein (vaccine strain and the consensus from the predominant Brazilian strains) and sialic acids.

### Epidemiological Profile of Influenza A and Statistical Analysis

The expected value of viral lineages per region of Brazil was determined based on the total frequency of each subclade across the country’s regions. This calculation involved multiplying the expected frequencies of each subclade by the total observed frequency for each region and subsequently dividing by the total number of observations within that region. We considered any difference outside the range of the critical value as significant. *χ*-square tests, with 0.05 as a significance level, were used to compare the proportions of viral lineages by macroregions of Brazil. All statistical tests were performed in R.

The sampling power of the combined dataset (GISAID and CeVIVAS) was calculated as described above, in the Clinical Sample Collection section

## Results

To better understand the panorama of the molecular epidemiology of Influenza circulating in Brazil between 2021 and 2023, we generated genome sequences of these viruses from different Brazilian States. We analyzed them together with Brazilian sequences available in the public database. Through analysis of the public data, the viral sequence data exhibited a steady rise over the last three years in Brazil. Still, the ratio of available HA gene sequences concerning positive cases did not exceed 12% (S1 Fig).

Our results revealed the circulation of A/H1N1pdm09, A/H3N2, and B/Victoria in Brazil between 2021 and 2023. It was possible to detect the predominance of the A/H3N2 virus in 2021/2022, and the co-circulation of the three viruses at the end of 2022. However, in 2023, our analyses indicated a sharp increase in the circulation of both B/Victoria and A/H1N1pdm09 viruses in the country (Fig 1).

**Fig 1.**
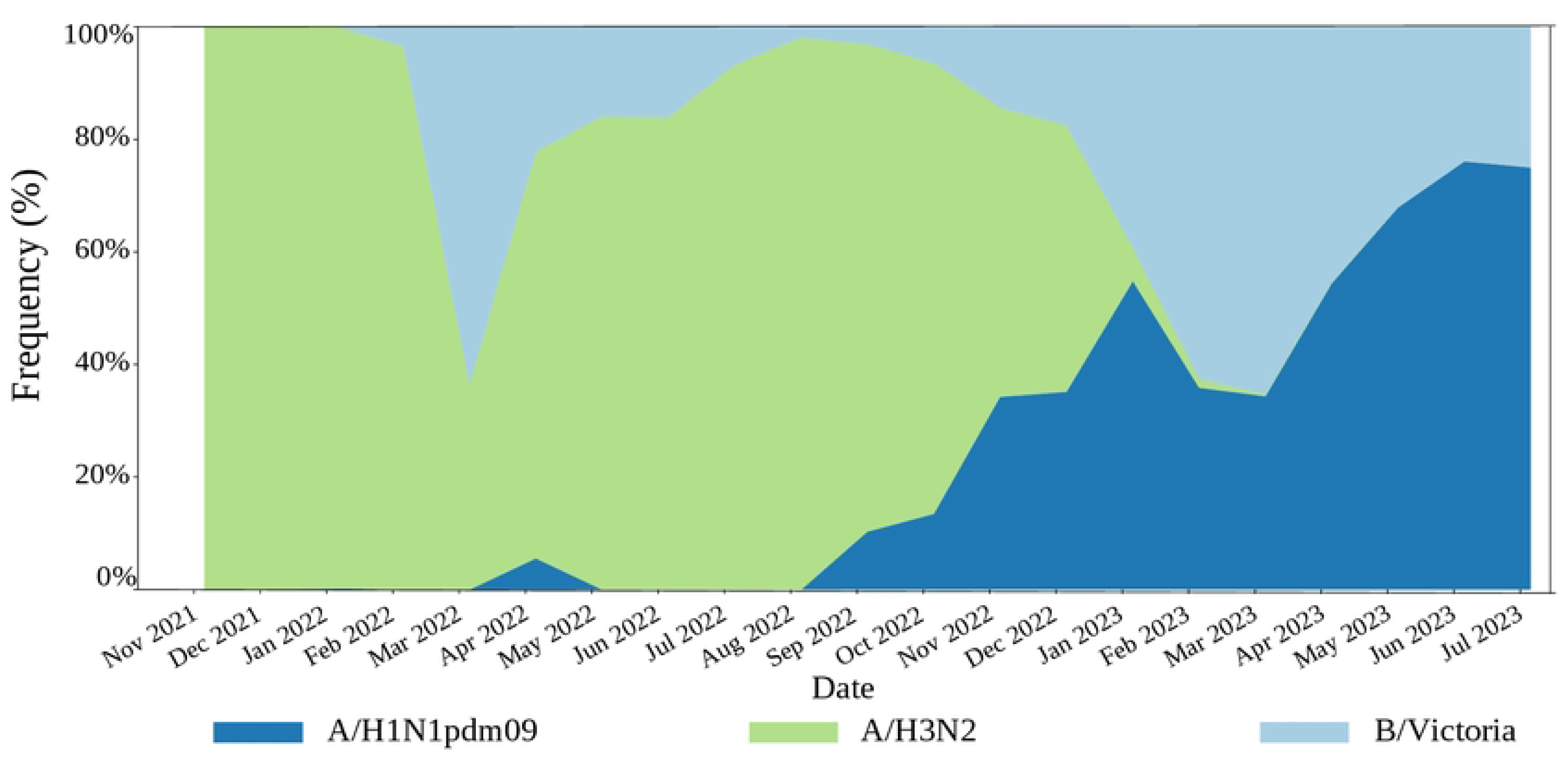
Frequency of Influenza viruses circulating in Brazil between 2021 and 2023 according to the performed phylogenetic analysis.

### Phylogenetic Analyses

A/H1N1pdm09 - Predominance of viruses belonging to the 5a.2a.1 subclade Phylogenetically, all 896 Brazilian sequences were grouped in three clade/subclades: almost in 5a.2a.1 (n=626, 70%) and, to a lesser extent, in subclade 5a.2a (n=252, 28%) and clade 5a.1 (n=18, 2%) (Fig 2A) that were circulating in the period when this study was concluded (first semester of 2023). In the 5a.2a.1 subclade, two main clusters were recovered. The first cluster (I-blue, n=37, 4% of the total) was characterized by T216A amino acid substitution and consisted of sequences primarily from the Southeast region (n=32). These sequences were also grouped as a sister group of the vaccine strain A/Victoria/4897/2022 - designated for administration in 2024 in the National Influenza Vaccination Campaign [48]. The second cluster (II-blue, n=592, 65.8% of the total) encompasses most Brazilian sequences from all regions of the country. From that specific cluster, 115 sequences exhibited the T270A amino acid substitution, with a notable prevalence in the North (n=61) and Northeast (n=34) regions (Fig 2B).

**Fig 2.**
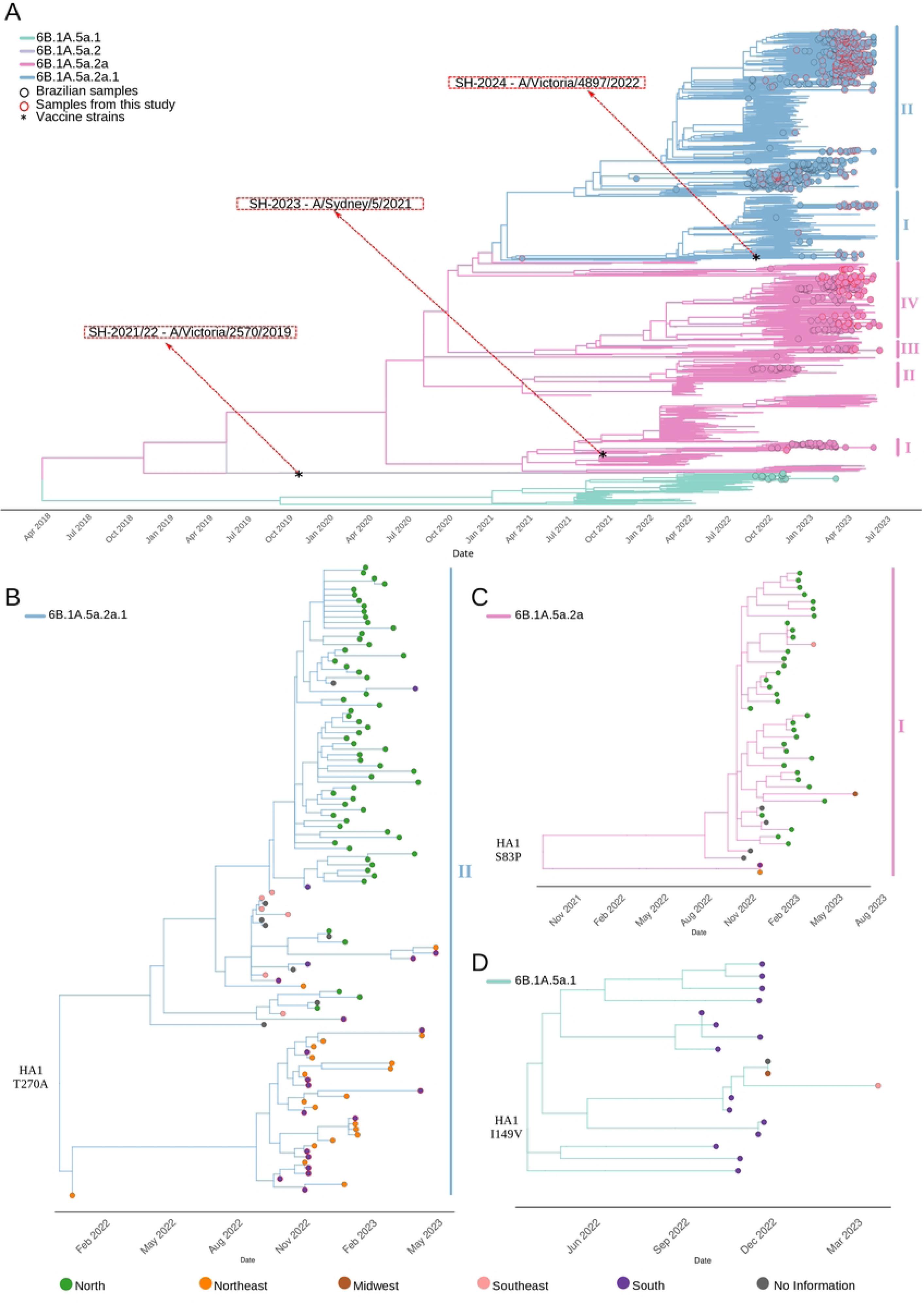
**Worldwide time-aware phylogeny of the genomic segment Hemagglutinin (HA) of the A/H1N1pdm09 viruses collected between January 2021 and July 2023 constructed by Nextstrain.**(A) In Brazil, subclades 5a.2a.1 (clusters I-II) and 5a.2a (clusters I-IV) are notably predominant. Southern Hemisphere recommended vaccine strains from 2021 to 2024 are indicated by asterisks and red arrows. The black circles identified Brazilian sequences; the red circles identified sequences generated in this study. (B) Zoom-out of 5a.2a.1 II cluster of Brazilian sequences mostly from North and Northeast regions with HA1 T270A amino acid substitutions. (C) Zoom-out of 5a.2a I cluster consisted of sequences mainly from the North region with HA1 S83P amino acid substitution. (D) Zoom-out of 5a.1 grouping sequences mainly from the South region, characterized by HA1 I149V amino acid substitution.

Within subclade 5a.2a, five clusters were recovered: the first cluster (I-pink; n=43, 4.8% of the total) mainly consisted of sequences from the Northern macroregion (n=35) of the country displaying the S83P amino acid substitution (Fig 2C). Also, this cluster was recovered as a sister group of the vaccine strain A/Sydney/5/2021 – administered in 2023 by the National Influenza Vaccination Campaign [9]. The second cluster (II-pink; n=19, 2.1% of the total) grouped sequences from the Southern (n=17) region and was characterized by A48P amino acid substitution. The third cluster (III-pink; n=13, 1.4% of the total) predominantly gathered sequences from Brazil’s Northeast (n=9) region. Lastly, the fourth cluster (IV-pink, n=170, 19% of the total), grouped sequences from all five macroregions of the country.

Within 5a.1, all Brazilian sequences (n=18, 2% of the total; Fig 2D) were grouped into a single clade, characterized by the shared I149V amino acid substitution. This clade predominantly consisted of sequences from the Southern region (n=15) and fewer representatives from the Midwest (n=1), and Southeast regions (n=1).

The comparison between the HA consensus of viruses circulating between 2021/2022 in Brazil and the vaccine strain egg-propagated A/Victoria/2570/2019 - vaccine administered in the 2021/2022 seasons, revealed 8 to 16 amino acid substitutions.

Compared with the predominant Brazilian subclade (5a.2a.1) in 2022, 14 amino acid substitutions were identified, 11 of which occurred in epitope regions. Notably, three substitutions (R223Q, D260E, T277A) stand out by the position at the receptor binding and Fusion domains according to protein structure prediction (Fig 3). In contrast, fewer amino acid substitutions (4-10) were observed when comparing the predominant subclades collected in 2023 in Brazil with the egg-propagated vaccine strain A/Sydney/5/2021. Four common amino acid substitutions (N94D, A216T, R223Q, and I418V) were identified upon comparison of the consensus sequences from subclades 5a.2a.1 and 5a.2a with the strain A/Sydney/5/2021. The R223Q mutation is specifically at the residue responsible for interaction with sialic acid. As presented in Fig 3A, none of those mutations were close to the region with interaction with some antibodies like CR 6261. For more details see S1 and S2 Tables.

**Fig 3.**
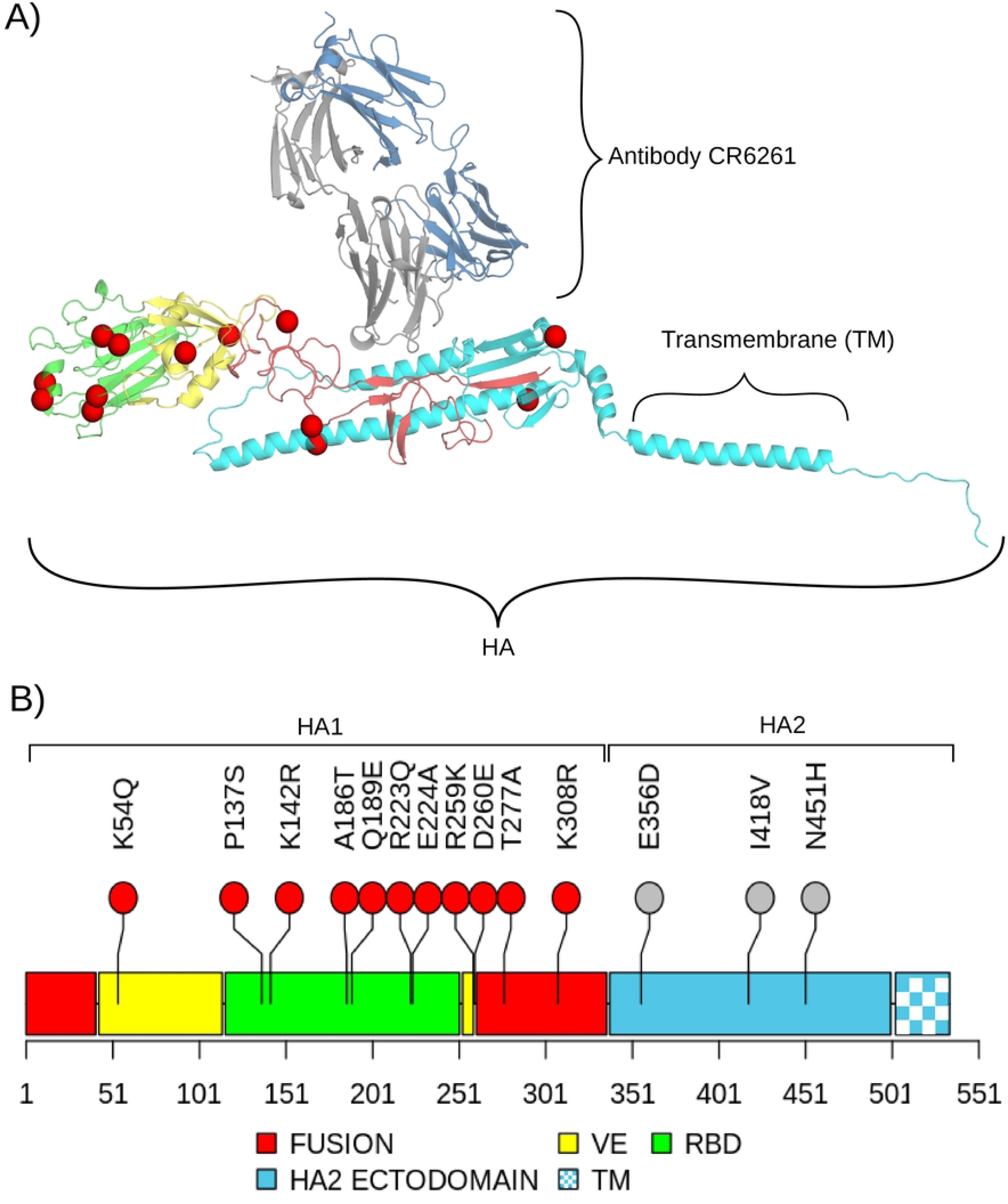
**Influenza virus A/H1N1pdm09 hemagglutinin (HA) protein structure and observed mutations between vaccine strain egg-propagated A/Victoria/2570/2019 and Brazilian consensus from subclade 6B.1A5a.2a.1 (5a.2a.1) in 2022.**(A) 3D predicted structure of complex antibody-HA, colored by different domains. Fusion domain (red), HA2 ectodomain (cyan); VE vestigial esterase domain (yellow), RBD receptor binding domain (green), TM anchor transmembrane anchor (checkerboard cyan); Substitutions in HA epitope regions indicated by red spheres. The antibody is colored by different chains: light chain (gray), and heavy chain (sky-blue). (B) 1D schematic view of HA domains, colored by different domains. Color coding as per panel (A). The main amino acid substitutions are shown in epitope positions (red circle), and non-epitope region (gray circle).

According to the virtual docking assay, the vaccine strain A/Victoria/2570/2019 RBD residues as H180, D187, K219, R223, E224, and T133 are responsible for hydrogen interactions with alpha-2,3 or alpha-2,6 sialic acids, while N130, K142, and D222 residues could be responsible for distance interactions. When examining the consensus sequence of subclade 5a.2a.1, we noted that only residues D187 and Q223 exhibited hydrogen interactions, while W150 potentially engaged in a CH-pi interaction with alpha-2,3 sialic acid. Regarding alpha-2,6 sialic acid, we observed that residue L191 could partake in a hydrophobic interaction with the substrate, while residues D187 and Q223 persisted in hydrogen interactions. For more detailed information see S2 Fig.

### Influenza A/H3N2 - High Diversification and Low Circulation during the 2023 Season

Phylogenetically, all 2,265 Brazilian sequences were clustered mainly within subclade 3C.2a1b.2a.2a.3 (2a.3, n=1687, 74.5%), followed by subclade 3C.2a1b.2a.2b (2b, n=456, 20.1%), and to a lesser extent in subclades 3C.2a1b.2a.2c (2c, n=76, 3.4%), 3C.2a1b.2a.2a (2a.2a, n=35, 1.5%), 3C.2a1b.2a.2a.1 (2a.1, n=9, 0.4%), and 3C.2a1b.2a.2a.1b (2a.1b, n=2, 0.1%) (Fig 4A).

**Fig 4.**
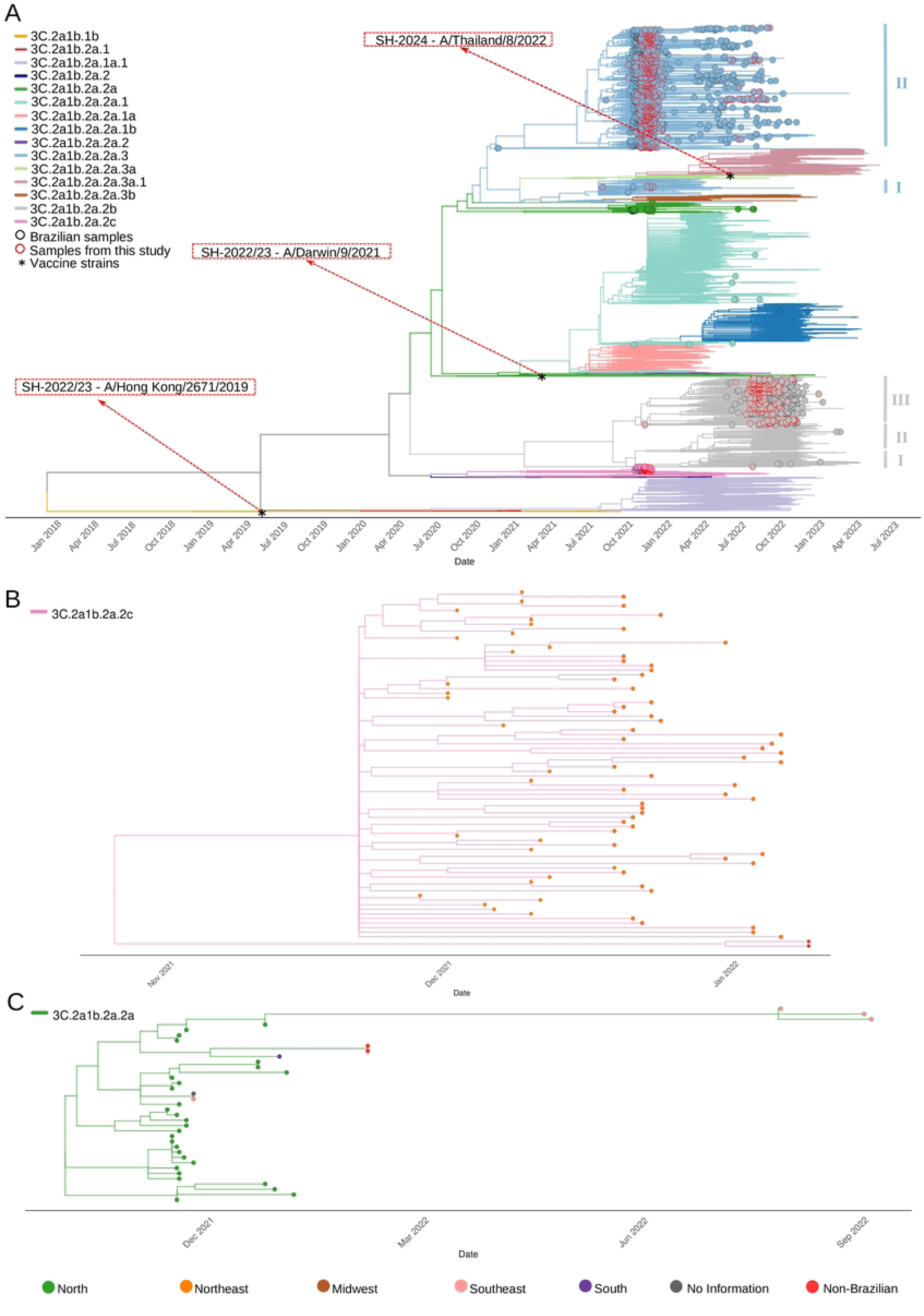
**Worldwide time-aware phylogeny of the genomic segment Hemagglutinin (HA) of the Influenza A/H3N2 collected between January 2021 and July 2023 constructed by Nextstrain.**(A) In Brazil, predominance and co-circulation of the subclades 2a.3 (clusters I-II blue), and 3C.2a1b.2a.2b (clusters I-III gray). (B) Zoom-out of subclade 2c, demonstrating exclusively Northeast sequences. (C) Zoom-out of subclade 2a.2a which was comprised mostly of sequences from the North region. The black circles identified Brazilian sequences; the red circles identified sequences generated in this study. Southern Hemisphere recommended vaccine strains from 2021 to 2024, which are indicated by asterisks and red arrows.

In our tree, within subclade 2a.3 two clusters were recovered: the first (I-blue, n=10, 0.4%) comprised only six sequences from the Midwest region, and the second cluster (II-blue, n=1677, 74%) encompassed most Brazilian sequences from all regions of the country. In subclade 2b, three clusters were recovered: the first cluster (I-gray, n=4, 0.2%) consisted of sequences originating from the South (n=3) and North (n=1) regions sharing the I242M amino acid substitution. In the second cluster (II-gray, n=2, 0.8%), the sequences from the South (n=2) presented S262N, T135A amino acid substitution. Within the third cluster (III-gray, n=449, 19.8%) most sequences originated from all regions of the country. The subclade 2c comprised exclusively of Northeast sequences (n=76, 3.3%, Fig 4B). Also, this subclade was recovered as the sister group of the 3a.1, which includes the vaccine strain Thailand/8/2022 - designated for administration in 2024 in the National Influenza Vaccination Campaign (BRASIL, 2024). In subclade 2a.2a, most of the sequences were from the North (n=35, 1.5%, Fig 4C). Among the subclade 2a.1, sequences from all regions except the North were grouped. Within the 2a.1b subclade, only sequences from the Northeast (n=2) were recovered.

The comparison between the HA consensus of viruses circulating during 2021 in Brazil and the vaccine strain egg-propagated A/Hong Kong/2671/2019, which constituted the vaccine administered in the 2021 season [22], revealed 18 to 23 amino acid substitutions. In contrast, fewer amino acid substitutions (5-8) were observed when comparing viruses collected in 2022 in Brazil with the vaccine strain egg-propagated A/Darwin/9/2021 - which was administered from April 2022 to May 2023 during the National Influenza Vaccination Campaign [49]. Upon comparing A/Darwin/9/2021 and the prevailing Brazilian subclade (2b) during the second semester of 2022, our analysis unveiled 6 amino acid substitutions, all localized within epitope regions (Fig 5). Mutations at residue 186 and 225 located within the sialic acid binding site were identified across all A/H3N2 virus lineages circulating in Brazil between 2021 and 2023.

**Fig 5.**
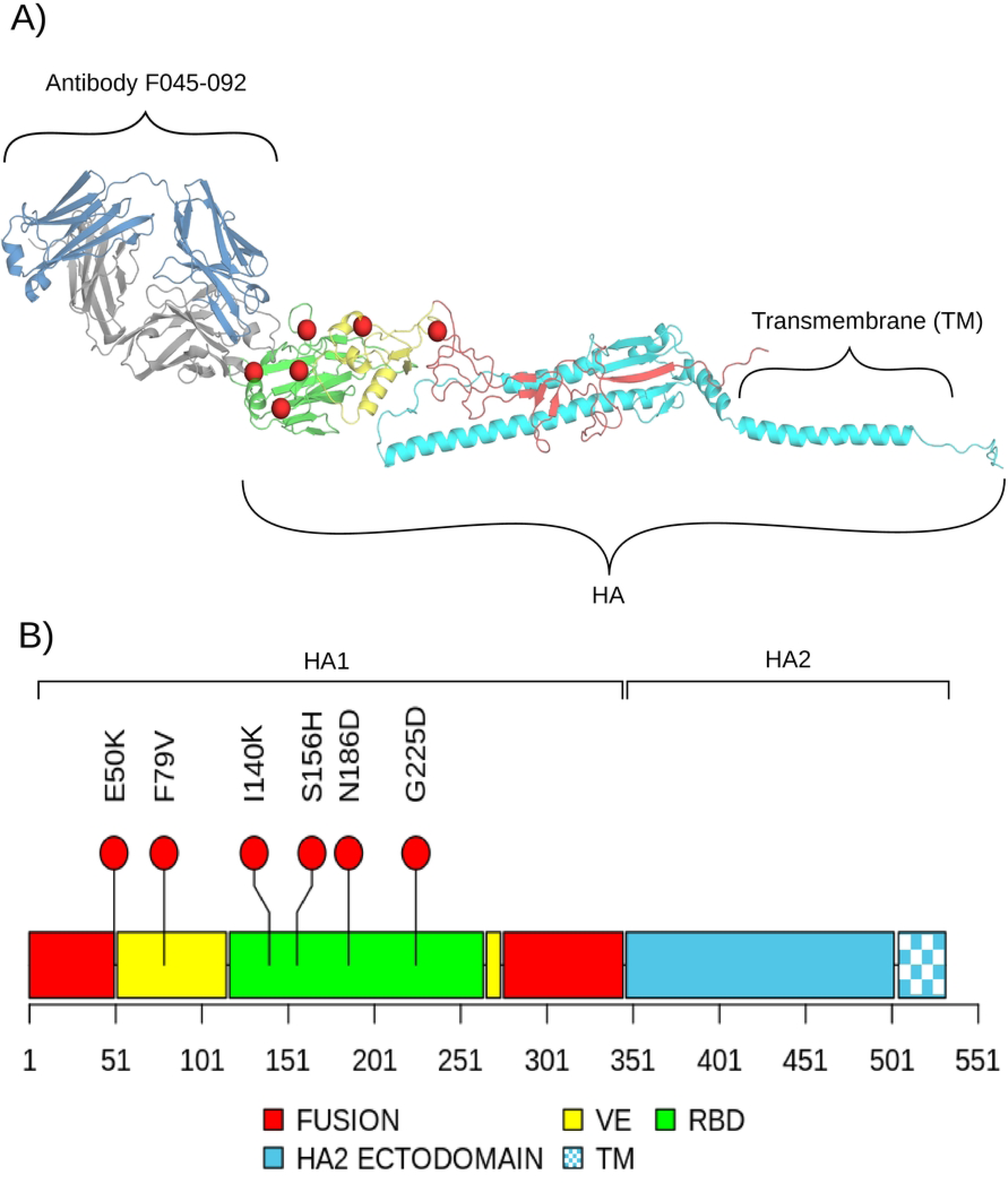
**Influenza virus A/H3N2 hemagglutinin (HA) protein structure and observed mutations between vaccine strain egg-propagated A/Darwin/9/2021 and Brazilian consensus from subclade 3C.2a1b.2a.2b (2b) in 2022.**(A) 3D predicted structure of complex antibody-HA, colored by different domains. Fusion domain (red), HA2 ectodomain (cyan); VE vestigial esterase domain (yellow), RBD receptor binding domain (green), TM anchor transmembrane anchor (checkerboard cyan); Substitutions in HA epitope regions indicated by red spheres. The antibody is colored by different chains: light chain (gray), and heavy chain (sky-blue). (B) 1D schematic view of HA domains, colored by different domains. Color coding as per panel (A). The main amino acid substitutions are shown in epitope positions (red circle).

This mutation resulted in a change of charge from an uncharged polar residue (Asparagine - N or Glycine - G) to a negatively charged residue (Aspartate - D). For more details, see S3 and S4 Tables. Electrostatic analyses indicated that E50K and F79V amino acid substitutions are in the fusion (F’) and vestigial esterase (VE) domains of HA1, respectively (Fig 5). The E50K substitution changes the electrostatic profile, while F79V could promote a change in the orientation of the side chain of amino acids close to it (S3 Fig). The I140K, N186D, and G225D mutations induce a little difference in the electrostatic profile of the RBD domain, making it a little more electro-negative to the RBD domain of the vaccine strain (S4 Fig). As presented in Fig 5A, all those mutations were close to the region with interaction with some antibodies like F045-092.

### Influenza B/Victoria-lineage - High Circulation in 2023 with Regional Arrangement

The obtained phylogenetic tree revealed that all 778 Brazilian Influenza B viruses were classified as Victoria lineage from the V1A.3a.2 clade (Fig 6A) and among those, almost presented the D197E amino acid substitution (n=649, 83.4%). Within the V1A.3a.2 clade were recovered nine clusters. The first (I, n=42, 5.4%) was composed mostly of sequences from the North (n=40) region of Brazil, in which T182A, D197E, and T221A were characterized (Fig 6B. The second cluster (II, n=85, 11%) encompasses sequences mostly from the Northeast, and a few sequences from all regions of the country (Fig 6C). This cluster was characterized by S208P, E128K, and A154E substitutions. The third (III) cluster encompassed two sequences from the Northeast and shared G141R substitution. Also, this cluster was recovered as a sister group of the recommended vaccine strain B/Austria/1359417/2021 – which made up the vaccine administered from 2022 and the 2024 seasons (BRASIL, 2024, 2023a, 2022). The fourth (IV) cluster comprised only one sequence from the Northeast region and displayed D197E, V117I, A154T, K326R, and 128K amino acid substitution. The fifth cluster (V, n=104, 13.4%), grouped sequences from all regions and can be characterized by D129G, E183K, and V87A substitutions. The sixth cluster (VI, n=83, 10.7%) grouped sequences (n=83) mostly from the Northeast (n=31) and North (n=30) regions, which shared the HA1 D197E, and Q200P amino acid substitutions (Fig 6AD). The seventh (VII) cluster contains most of the Brazilian sequences (n=315), of which almost all come from the Southeast and Northeast regions, both represented in nearly equal proportions. In the eighth cluster (VIII, n=10, 1.3%) were grouped sequences from all Brazilian regions, except the South, and presented D197E, and E183K substitutions. In the ninth cluster (VIII, n=107, 13.7%), almost all sequences were from the Northeast (n=97) region, and some were from the Southeast (n=8) and North (n=2) regions. This cluster was characterized only by D197E amino acid substitution.

**Fig 6.**
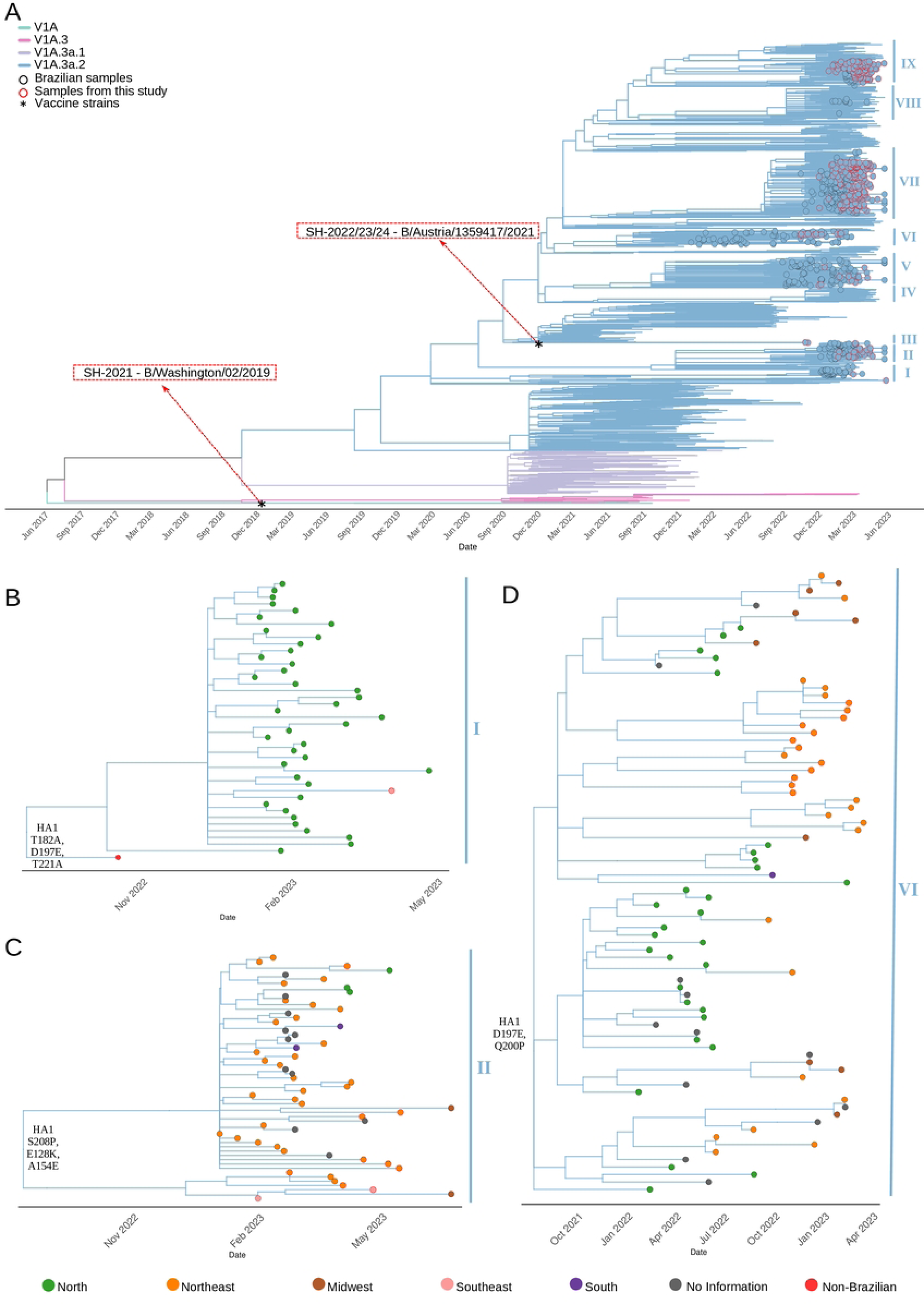
Worldwide time-aware phylogeny of the genomic segment Hemagglutinin (HA) of the Influenza B/Victoria-lineage viruses collected between January 2021 and July 2023 constructed by Nextstrain. (A) In Brazil, it was exclusively detected the circulation of the V1A.3a2 subclade. (B) Zoom-out of cluster I (characterized by HA1 T182A, D197E, and T221A amino acid substitutions), and composed mostly of sequences from the North region. (C) Zoom-out of cluster II comprises mostly sequences from the Northeast, with S208P, E128K, and A154E amino acid substitutions. (D) Zoom-out of cluster VI, grouping sequences mostly from the Northeast and North regions, sharing HA1 D197E, and Q200P amino acid substitutions. The black circles identify Brazilian sequences; the red circles identify sequences generated in this study. A red circle identifies the sequences generated in this study. The Southern Hemisphere recommended vaccine strains from 2021 to 2024 are indicated by asterisks and red arrows.

The comparison between the HA consensus of the viruses circulating during 2022 with the vaccine strain egg-propagated B/Austria/1359417/2021, revealed only two amino acid substitutions (D197E, and Q200P). In contrast, only one (D197E) amino acid substitution was observed when comparing viruses collected in Brazil in 2023 (S5 Table).

### Epidemiological profile of Influenza A viruses in macroregions of Brazil

Our phylogenetic analysis suggested that there are variations in Influenza lineage distribution across the Brazilian territory. Since Brazil contains a diversity of climate and demographic characteristics across the five macroregions, we decided to check if there are different circulation patterns across different regions. Therefore, we first checked if we had enough data to perform this comparison. Indeed, the number of sequences analyzed of A/H3N2 virus predominated between November 2021 and November 2022 provided a sample power of 98% (CI 97%), whereas the A/H1N1pdm09 sequences predominated between November 2022 and May 2023 provided a sample power of 87% (CI 90%). Following this, we conducted a subsampling of the final dataset, specifically targeting peak periods of predominance for the A/H1N1pdm09 and A/H3N2 subtypes, aiming to elucidate the nationwide epidemiological profile of Influenza A. Concerning A/H3N2 lineages (Fig 7), we observed a significantly higher frequency of 2a.3 in the Midwest region compared to other regions (chi-square, p*<*0.05), while subclade 2b exhibited higher circulation (chi-square, p*<*0.05), particularly in the Southeast. Additionally, the 2a subclade was only detected in three regions, with the North region registering significantly higher circulation than expected (chi-square, p*<*0.05).

**Fig 7.**
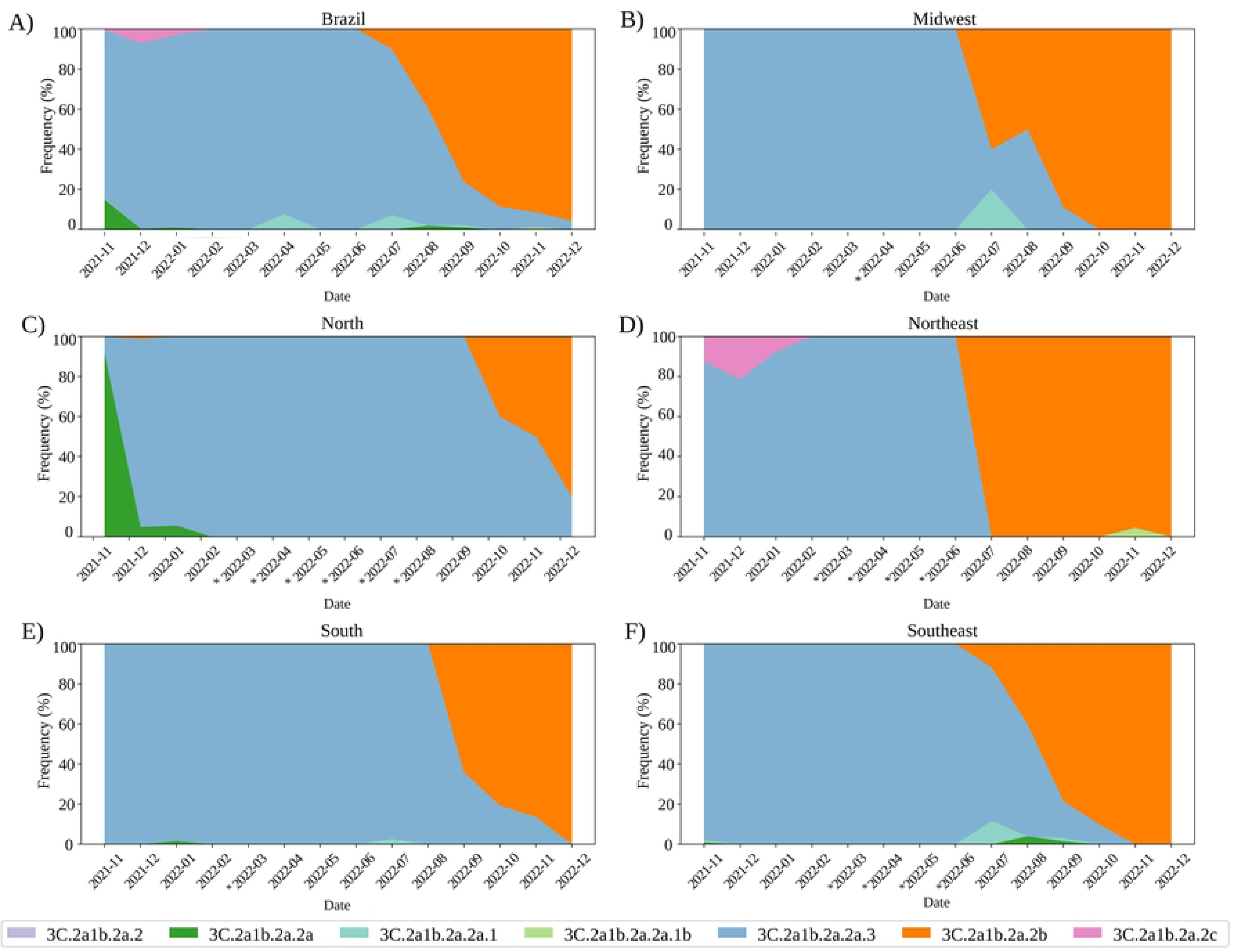
Frequency of A/H3N2 subclades 3C.2a1b.2a.2a.1 (2a.1), 3C.2a1b.2a.2a.3 (2a.3), 3C.2a1b.2a.2a.2b (2b), 3C.2a1b.2a.2 (2), 3C.2a1b.2a.2a (2a), and 3C.2a1b.2a.2c (2c) circulating from November 2021 to November 2022 in Brazil. (A) Frequency of A/H3N2 subclades in Brazil. (B) Frequency of A/H3N2 subclades in Midwest. (C) Frequency of A/H3N2 subclades in North. (D) Frequency of A/H3N2 subclades in Northeast. (E) Frequency of A/H3N2 subclades in South. (F) Frequency of A/H3N2 subclades in Southeast. (*) period of absence from genomic monitoring.

Meanwhile, viruses from the 2c subclade were exclusively recorded within the Northeast region. For detailed statistical information, see S6 Table. In respect of A/H1N1pdm09 viruses (Fig 8), the prevalence of 5a.2a.1 was notable across all macroregions of the country, except in the North and South regions, where this strain circulated at lower levels (chi-square, p*<*0.05), while, in the northern region, a higher circulation of 5a.2a strain was detected. Despite the low frequency of the 5a.1 strain at the national level, a higher circulation was observed in the Northeast and southern regions. For detailed statistical information, see S7 Table.

**Fig 8.**
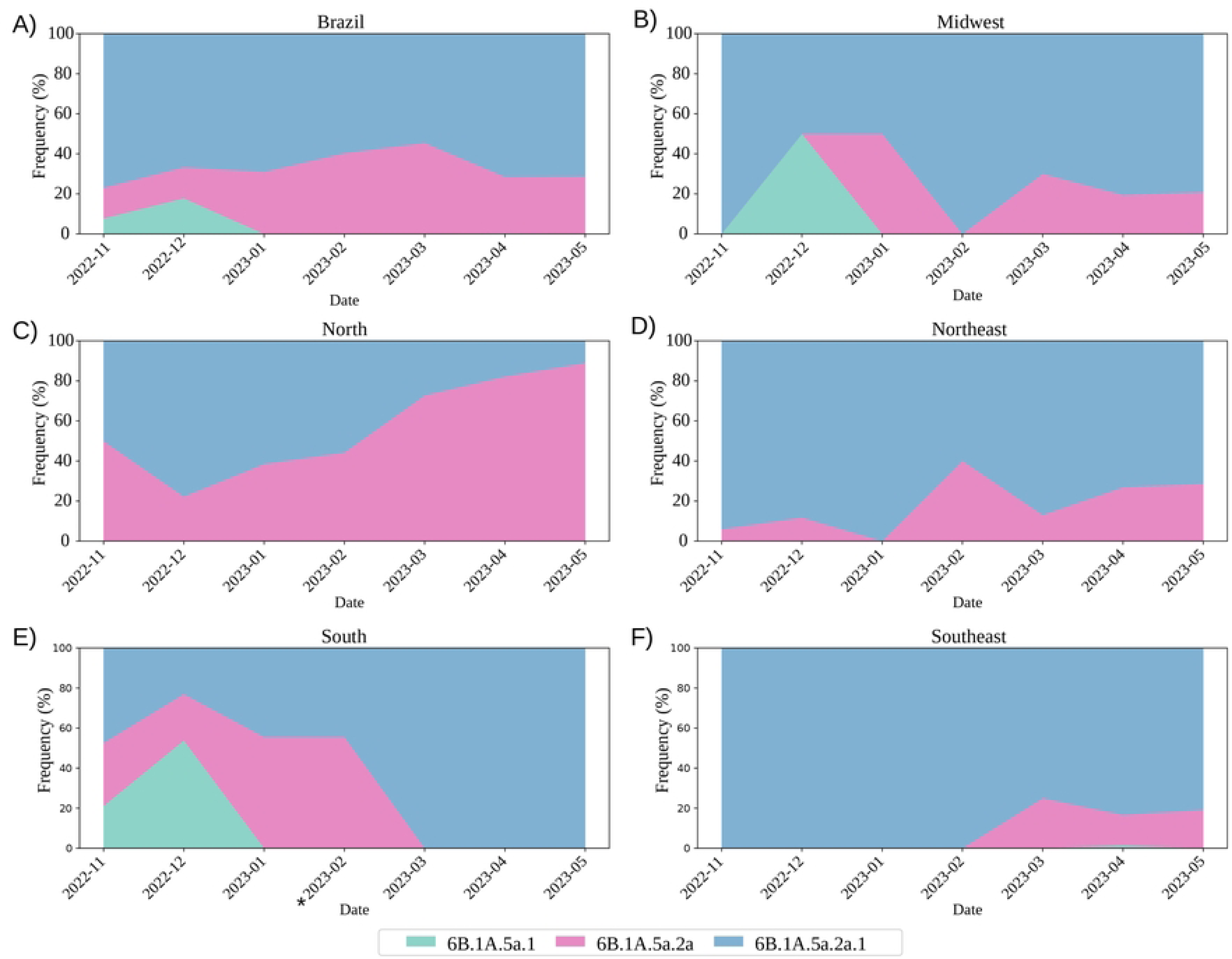
Frequency of A/H1N1pdm09 subclades 6B.1A.5a.1 (5a.1), 6B.1A.5a.2a (5a.2a) and 6B.1A.5a.2a.1 (5a.2a.1) from November 2022 to May 2023 in Brazil.(A) Circulation in Brazil; (B) Frequency of A/H1N1pdm09 subclades in Midwest. (C) Frequency of A/H1N1pdm09 subclades in North. (D) Frequency of A/H1N1pdm09 subclades in Northeast. (E) Frequency of A/H1N1pdm09 subclades in South. (F) Frequency of A/H1N1pdm09 subclades in Southeast. (*) period of genomic monitoring absence.

## Discussion

The number of positive Influenza cases in Brazil increased from 5,252 in 2021 to 18,736 in 2023, indicating a resurgence in viral activity compared to the pre-pandemic scenario (BRASIL, 2023a, 2023b, 2022). Notably, between September 2021 and January 2022, Influenza viruses circulated at significantly reduced rates, comprising less than 3% of cases, compared to the pre-pandemic rate of 17% [50]. The decrease in Influenza circulation has been linked to the widespread disruption caused by the COVID-19 pandemic [51]. This global health crisis prompted a concerted effort to enhance molecular surveillance of pathogen genomes, emerging as a critical measure for preparedness against future epidemics [52, 53]. The progressive sequencing efforts spanning from 2021 to 2023 in Brazil reflect this effort and have unveiled the identification of three Influenza viruses circulating in Brazilian territory: A/H1N1pdm09, A/H3N2, and B/Victoria lineage. This finding is consistent with data from the Brazilian Epidemiological Surveillance [22, 49, 54] and further supports the global co-circulation of Influenza A and B viruses.

Our phylogenetic analysis of A/H1N1pdm09 revealed a co-circulation of the 5a.2a.1, 5a.2a, and 5a.1 subclades. The predominance of 5a.2a.1 viruses in the country was consistent with observations made in other regions, such as North America and Europe [55]. The HA1 T216A amino acid substitution was observed in some sequences from the Southeast and was previously identified in many 5a.2a.1 viruses [37]. The T270A amino acid substitution observed in sequences from the North and Northeast regions was also observed as recurrent substitutions within the subclade. The functional significance of its substitution remains unknown [56, 57]. Among the five clusters recovered within 5a.2a, three of them were associated with the macroregions suggesting multiple localized introduction events. The cluster formed by sequences from the northern region which were characterized by the presence of the S83P substitution has also been identified in sequences from Thailand [58], the Middle East, Asia, and Oceania (as available at https://nextstrain.org/flu/seasonal/h1n1pdm/ha/2y?branchLabel=aa). This finding contrasts with previous studies that have suggested the Southeastern region of Brazil as the primary epicenter of H1N1pdm09 transmission, leading to widespread epidemics [59, 60] highlighting that other Brazilian regions may also act as a route for Influenza strains introduction.

The phylogenetic relationship recovered a closer relationship between the A/H1N1pdm09 Brazilian viruses and the vaccine strains. Additionally, the reduction in amino acid substitutions from 14 to 10 among the circulating viruses in 2023 compared to the vaccine strain A/Sydney/5/2021 suggests a low antigenic drift. Among the amino acid substitutions at antigenic sites, P137S, K142R, D260E, and T277A substitutions are characteristics of viruses belonging to the 5a.2a.1 subclade [39]. These substitutions were related to reduced geometric mean titers (GMTs) post-vaccination, especially to the egg-propagated vaccine strain A/Sydney/5/2021 [37]. The current Southern Hemisphere vaccine A/Victoria/4897/2022 covers the circulating clades [37].

A previous study highlights the complex evolution of A/H3N2, characterized by multiple clades globally distributed, posing challenges for vaccine composition [61]. Our analysis identified seven clades/subclades circulating in Brazil, with 2a.3 and 2b subclades being more prevalent. Within the 2a.3 subclade, Brazilian sequences grouped into two clusters, while the 2b subclade exhibited three clusters, one of which comprised 99% of sequences from all regions of Brazil. Despite the low circulation of viruses from the remaining subclades 2a.2a, 2c, and 2a.1b, they were restricted to the North and Northeast regions. The diversity of A/H3N2 viruses elucidated through our analysis corroborates the findings outlined in the World Health Organization (WHO) reports spanning the years 2021 to 2022 [38, 39, 50, 62]. These findings align with WHO reports and underscore H3’s rapid evolutionary rate [63, 64].

A comparison between the vaccine strain A/Hong Kong/2671/2019 (season 2021) and prevalent 2021 viruses from the 2a.3 subclade revealed 21 amino acid substitutions. Seventeen of these mutations are located within major antigenic sites, including E53N, K135T, and H156S. Notably, some of these substitutions (H156S, D53N) are characteristic of the 2a.3 subclade (WHO, 2023b). The discrepancy between circulating viruses and the 2021 vaccine strain prompted the update of the H3N2 component for the following season [38], resulting in fewer amino acid substitutions (5-7) in the predominant subclades in 2022. One (G225D) of the mutations is shared among the dominant subclades, and four others (E50K, F79V, I140K, and G186D) are typically encoded by viruses from the 2b subclade [55]. Notably, the G225D, I140K, and N186D are in the sialic acid binding site, and the E50K and F79V substitutions are situated in the F’ and VE domains [20, 65–67]. Mutations within the receptor-binding domain can alter the interaction between the HA protein and alpha-2,3 or alpha-2,6 sialic acids across diverse host species [68]. Despite its general conservation, the receptor-binding site is prone to selective pressures, novel mutations, and antibody evasion [69]. The G225D substitution resulted in a charge change from an uncharged polar residue, potentially allowing circulating strains to evade recognition by vaccine-induced antibodies. The F’ domain, located within the stem domain and implicated as a site of antibody binding, facilitates low pH-triggered membrane fusion activity of HA within endosomal compartments [20, 65, 70, 71]. The amino acid substitutions in the F’ domain, particularly those altering its electrostatic profile at lower pH levels, play a pivotal role in modifying viral infection dynamics [65]. Regarding VE domain mutations, while its function remains unclear for Influenza A and B viruses, it is known to be crucial for interaction with monoclonal antibodies, and alterations in this region could impact viral antigenicity [72]. Indeed, the post-infection ferret antisera raised against the egg-propagated A/Darwin/9/2021-like viruses exhibited a less favorable reaction, especially towards viruses expressing 2a.3a.1 or 2b HA genes [37]. Conversely, ferret antisera raised against 2a.3a.1-like viruses (e.g. egg-propagated A/Thailand/8/2022, season SH 2024) demonstrated robust recognition of most circulating viruses [37], and suggests a closer similarity between the circulating strains and the vaccine strain.

Considering the panorama of the Influenza B virus in Brazil, literature on the subject is limited. Therefore, our investigation represents an essential study in advancing our understanding of this virus in the country. Our analysis revealed a single main clade of Influenza B, identified as V1A.3a.2, has circulated in Brazil. This clade has been exclusively circulating not only in Brazil but also in other regions across the globe since February 2023, with a noteworthy substitution of the Yamagata lineage [73–76]. Our findings are consistent with previous studies conducted in the North and Northeast regions of Brazil, highlighting the unique dominance of the V1A.3a.2 clade in these areas [77]. The ongoing diversification of the clade has resulted in the emergence and near fixation of distinct subclades, some of which exhibit recurrent HA1:197E and HA1:183K substitutions [78, 79]. Recent viruses carrying the D197E mutation have been preliminarily classified as the C.5 clade, with seven descending major subclades (C.5.1 - C.5.7) identified globally [78]. In this study, four subclades (C.5.1, C.5.2, C.5.3, C.5.4) were identified as circulating in Brazil. Most sequences were recovered in C.5, followed by subclades C.5.3, and C.5.2. According to our phylogenetic analysis, clusters VII and IX are likely to be assigned to clade C.5, while clusters V and VI may be designated as C.5.3 and C.5.2, respectively. Both haplotypes C.5.2 and C.5.3 have been predominantly recorded in South America, suggesting their probable origin in this region (available at: https://nextstrain.org/flu/seasonal/vic/ha/2y, accessed on 5 April 2024).

The amino acid substitutions D197E and Q200P were identified in the consensus sequences of circulating Brazilian viruses in 2022. However, only the D197E substitution was observed in 2023. The WHO has previously documented the D197E mutation, and its presence did not suggest a decrease in vaccine efficacy [37]. This finding aligns with integrated hemagglutination inhibition (HI) data and molecular evolution analysis of the HA segment, indicating effective coverage against recent 3a.2 strains by the current vaccine strain B/Austria/1359417/2021 [79].

The analysis of viral alternation patterns revealed in this study provides a detailed insight into the circulation of Influenza viruses in Brazil. This dynamic closely aligns with the epidemiological data provided by the Brazilian Ministry of Health for the specified period [9, 49]). A similar trend of low A/H3N2 HA sequence counts, and an increase of both A/H1N1pdm09 and B/Victoria circulation was observed across different global regions throughout 2022, including Europe, North America, and Oceania [62, 80, 81]. Furthermore, between February and September 2023, a notable increase in the detection of the B/Victoria virus, from 5.8% to approximately one-third of globally detected viruses, was observed [37, 55].

In summary, our investigation unveiled significant findings, including the detection of clades exhibiting mutations suggestive of antigenic drift, heightened diversity within the A/H3N2 subtype, and distinct distribution patterns across different regions of Brazil. This research highlighted the critical importance of monitoring and understanding the molecular epidemiology of Influenza viruses. Such insights are instrumental in guiding public health agencies to implement effective measures for mitigating transmission risks and controlling outbreaks. Furthermore, the knowledge gained from this study can contribute to the development of vaccine strategies for a continental country.

## Supporting information

**S1 Fig. Positive cases of subtyped Influenza and available/deposited Hemagglutinin (HA) gene sequences between 2021 and 2023.** Column values refer to the left y-axis. The black line refers to the values in the right y-axis, which indicates the proportion of available/deposited HA gene sequences over the total number of positive subtyped Influenza cases in Brazil. Data was obtained on 27 December 2023 at https://platform.epicov.org. (*) data collected until the 30th epidemiological week.

**S1 Table. Amino acid substitution in the Hemagglutinin (HA).** Comparison between the consensus of the circulating A/H1N1pdm09 viruses from Brazil between 2021 and 2022 and Southern Hemisphere 2021-2022 vaccine strain egg-propagated A/Victoria/2570/2019.

**S2 Table Amino acid substitution in the Hemagglutinin (HA).** Comparison between the consensus of the circulating A/H1N1pdm09 viruses from Brazil 2023 and Southern Hemisphere 2023 vaccine strain egg-propagated A/Sydney/5/2021.

**S2 Fig. H1N1 docking analysis results.** In (A) and (B) are represented H1N1 interactions with alpha-2,3 sialic acid for A/Victoria/2570/2019 and 5a.2a.1 Brazilian consensus, respectively. The RDB domain is represented as cartoon (green), the main residues to interaction are represented as licorice, and alpha-2,3 sialic acid is represented as stick (gray). All interactions are represented in yellow dashes. In (C) and (D) are represented H1N1 interactions with alpha-2,6 sialic acid for A/Victoria/2570/2019 and 5a.2a.1 Brazilian consensus, respectively. The RDB domain is represented as cartoon (green), the main residues to interaction are represented as licorice, and alpha-2,3 sialic acid is represented as stick (gray). All interactions are represented in yellow dashes. The results show that 5a.2a.1 Brazilian consensus has more affinity to alpha-2,6 sialic acid than alpha-2,3 sialic acid while A/Victoria/2570/2019 has a similar affinity to both sialic acids.

**S3 Table. Amino acid substitution in the Hemagglutinin (HA).** Comparison between the consensus of the circulating A/H3N2 viruses collected in Brazil in 2021 and Southern Hemisphere 2021 vaccine strain egg-propagated A/Hong Kong/2671/2019.

**S4 Table. Amino acid substitution in the Hemagglutinin (HA).** Comparison between the consensus of the circulating A/H3N2 viruses collected in Brazil between 2022 and 2023 compared with the Southern Hemisphere 2022-2023 vaccine strain egg-propagated A/Darwin/9/2021.

**S3 Fig. Electrostatic analysis of amino acid changes on the VE domain from A/H3N2.** In (A) and (B) We observed the protein electrostatic profile for egg-propagated vaccine strain A/Darwin/9/2021 and 3C.2a1b.2a.2b (2b) subclade, respectively. In (C) and (D) are indicated a zoom in the VE domain from egg-propagated vaccine strain A/Darwin/9/2021 and 3C.2a1b.2a.2b (2b) subclade, respectively. The main electrostatic changes caused by E50K and F79V can be observed in (C) and (D) figures.

**S4 Fig. Electrostatic analysis of amino acid changes on the RBD domain from A/H3N2.** In A) and B) We observed the protein electrostatic profile for egg-propagated vaccine strain A/Darwin/9/2021 and 3C.2a1b.2a.2b (2b) subclade, respectively. In (C) and (D) are indicated a zoom in the VE domain from egg-propagated vaccine strain A/Darwin/9/2021 and 3C.2a1b.2a.2b (2b) subclade, respectively. The main electrostatic changes caused by I140K, N186D, and G225D mutations on the RBD domain can be observed in (C) and (D) figures.

**S5 Table. Table. Amino acid substitution in the Hemagglutinin (HA).**Comparison between the consensus of the circulating B/Victoria-lineage viruses collected in Brazil between 2022 and 2023 and Southern Hemisphere 2022-2023 vaccine strain egg-propagated B/Austria/1359417/2021.

**S6 Table. Adjusted deviation from expectation and sample size.** Values of A/H3N2 lineages circulating in Brazilian macroregions from November 2021 to December 2022.

**S7 Table. Adjusted deviation from expectation and sample size.** Values of A/H1N1pdm09 lineages circulating in Brazilian macroregions from November 2022 to May 2023.

**S1 File. Supporting Information for Material and Methods.** This document supports Clinical Sample collection, Ethics approval and consent to participate, Genomic library preparation and Influenza next-generation sequencing, and Genome Assembly Pipeline.

**S8 Table. Final dataset used in this study.** This file includes 3,966, 8,609, and 3,996 sequences from A/H1N1pdm09, H3N2, and B/Victoria-lineage, respectively. All sequences generated in this study were deposited in the GISAID database.

## Acknowledgments

The authors are grateful to Durval de Moraes Junior and Claudia Anania Santos da Silva for respectively logistic and information technology support.

